# Thermodynamics and selection of the plasminogen activator inhibitor-1 latency transition

**DOI:** 10.1101/2025.06.03.655624

**Authors:** Laura M Haynes, Matthew L Holding, Jaie Woodard, David Siemieniak, David Ginsburg

## Abstract

Plasminogen activator inhibitor-1 (PAI-1), like other serine protease inhibitors (SERPINs), exists in a functionally active metastable conformation. Unlike other SERPINs, PAI-1 undergoes a relatively rapid rearrangement to a lower energy latent conformation. We employ deep mutational scanning (DMS) to simultaneously probe the effects of nearly all single amino acid substitutions (92%) in PAI-1 on its latency transition as well as its ability to inhibit its canonical protease target, urokinase-type plasminogen activator (uPA). The DMS results are interpreted in the context of variant effect predictors (EVE, AlphaMissense, and CPT) and protein stability predictors (EvoEF and FoldX), as well as the evolutionary conservation of the PAI-1 sequence space across extant vertebrate species. We demonstrate that while variant effect predictors can generally partition PAI-1 as functional or non-functional with respect to uPA inhibition, they perform poorly when attempting to discriminate the effects of PAI-1 variants on its latency transition. However, by employing protein stability predictors, we demonstrate that the PAI-1 latency transition is most likely driven by changes in the energy barrier to the latency transition. Finally, we show that PAI-1’s ability to inhibit uPA, as well as its thermodynamic instability in its active conformation are both under purifying selection that limits the sequence space available to PAI-1. These findings suggest that DMS of collateral fitness effects, including PAI-1’s latency transition, may be better interpreted in contexts other than variant effect predictors, including protein thermodynamics and evolution.

## INTRODUCTION

Plasminogen activator inhibitor-1 (PAI-1) is a member of the serine protease inhibitor (SERPIN) protein superfamily (1, 2). It is a key regulator of fibrinolysis, or the enzymatic breakdown of fibrin clots, through its inhibition of urokinase- and tissue-type plasminogen activators (uPA and tPA). High levels of PAI-1 have been associated with numerous pathophysiologies (1, 3–5); however, the precise role and function of PAI-1 in many of these pathways have yet to be elucidated. Complete PAI-1 deficiency is associated with a mild-to-moderate bleeding diathesis (6, 7). Conversely, PAI-1 haploinsufficiency has been linked to increased longevity (8).

PAI-1, like other inhibitory SERPINs, exists in a metastable active conformation in which the reactive center loop (RCL), extends from the core of the protein (**Fig. 1**) and contains an amino acid sequence that mimics that of the target protease’s preferred cleavage site (9). When the protease attempts to cleave the RCL’s scissile bond, it becomes trapped as the RCL inserts into the SERPIN’s β-sheet A prior to hydrolysis of the acyl intermediate. Thus, in the *inhibitory pathway*, the SERPIN and serine protease become covalently linked in a lower energy conformation that renders both no longer functional. PAI-1 may also exist in other low energy conformations under two additional circumstances: (i) *the substrate pathway*—when hydrolysis of the acyl intermediary occurs before insertion of the RCL into β-sheet A, or (ii) a *latency transition*—when the RCL spontaneously inserts partially into β-sheet A in the absence of a proteolytic event. Among SERPINs, the latter is a unique feature of PAI-1 as it undergoes this latency transition with a relatively short half-life (t_1/2_ = 1-2 h) (10–13).

**Figure 1.**
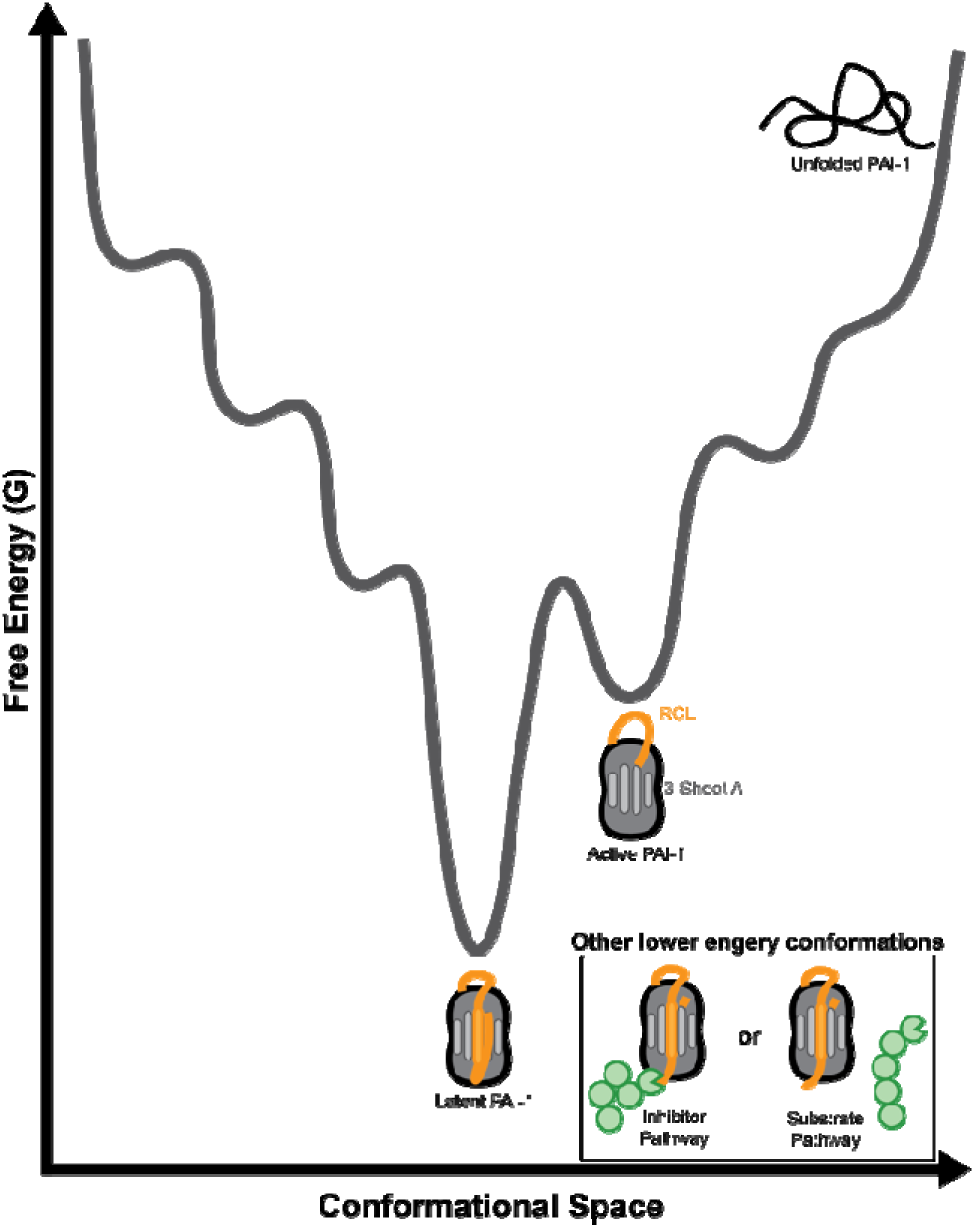
Functional PAI-1 exists as a metastable protein in a local energy minimum. The protein folding energy landscape is shown as a function of both conformational space (*x-axis*) and free energy (G, *y-axis*). Unfolded PAI-1 exists in the highest energy state with the most accessible conformations. Active PAI-1 occupies a local energy minimum in which the RCL (*orange*) extends from the globular core of the protein. To transition to its inactive, lowest energy latent conformation, PAI-1 must overcome an energy barrier that corresponds to the insertion of the RCL into β-sheet A (*light grey*). PAI-1 also enters additional lower energy conformations following a reaction with a target protease (*green*) through either the inhibitor or substrate pathway that both result in RCL insertion into β-sheet A.

The vast majority of proteins exist in their lowest energy conformation in which Gibbs free energy of folding (ΔG) between the unfolded and the functional native state is the most negative. However, metastable proteins, including SERPINs, exist in a functional conformational state that occupies a local energy minimum on its protein folding energy landscape (**Fig. 1**) (14). SERPINs take advantage of the energetic differences between their functional conformation and their reacted states (inhibitor or substrate pathways) as their inhibitory function is driven by exergonic conformational changes (ΔG<0). During its latency transition, as described above, PAI-1 collapses to a lower energy conformation in the absence of a proteolytic challenge; however, other SERPINs do not undergo this relatively rapid transition despite similarities in structure. Based on recombinant expression of shark and lungfish PAI-1, the latency transition appears to be conserved across species (15). However, we are only just beginning to understand the contributions of individual amino acid residues to this property (16–19).

Variant effect predictors (VEPs) offer the promise of being able to interpret the effects of protein sequence variation in lieu of DMS studies and, thereby identify potentially actionable pathogenic variants in patients (20–23). However, VEPs are often unable to identify more subtle phenotypic changes in protein function (24). An alternative and complementary approach to VEPs is protein stability predictors (PSPs), which are computational algorithms that estimate the relative change in the Gibbs free energy of folding (ΔΔG) resulting from a given amino acid substitution using structural information (25, 26). As the lowest energy conformation represents the functional conformation for most proteins, lower ΔΔG values represent more stable amino acid substitutions, while larger ΔΔG values correspond to destabilizing amino acid substitutions.

Therefore, given PAI-1’s metastable nature, we hypothesized that comparing effects of amino acid substitutions on ΔΔG in its active and latent conformations might provide insight into the thermodynamics of this transition that cannot be assessed by VEPs.

We have previously used deep mutational scanning (DMS) to map PAI-1’s mutational landscape, characterizing substitutions that impact both uPA inhibition (27, 28) and the timing of the latency transition (17). Here, with the addition of our improved coverage of PAI-1’s available sequence space, we expand upon these findings in three ways. First, we compare the results of our DMS screen to VEPs for predicting missense mutation pathogenicity. Second, we relate the DMS screen to PSPs for evaluating PAI-1’s canonical function as a uPA inhibitor and for its latency transition. Third we examine PAI-1’s natural sequence variation in the context of amino acid subsititions identified in our DMS screen as impacting both PAI-1’s inhibitor function and latency transition. We approach these questions using an integrative approach that includes developing a site saturation variant (SSV) library that has nearly equal representation of all single amino acid substitutions in PAI-1, allowing us to refine and expand upon our previous studies (17, 27, 28). To analyze the SSV screen, we incorporate approaches from computational, structural, and evolutionary comparative biology. Combined, these approaches provide insights into the thermodynamics governing PAI-1’s latency transition and demonstrate that both PAI-1’s function as a uPA inhibitor, as well as its relatively rapid latency transition, are features that dictate the amino acid sequence space available for PAI-1 evolution.

## RESULTS

### Effects of single amino acid substitutions on PAI-1 function and stability

The PAI-1 site-saturation variant (SVV) library used here contains 6,879 single amino acid substitutions (96% of the 7,201 total potential single amino acid substitutions) with approximately equal representation of each variant. We selected for variants that inhibited uPA before (0h) or after a 48h incubation at 37°C (**Fig. 2A-C; SI Data 1**). We then used principal component analysis (PCA) to visualize the between-sample variation in amino acid substitution enrichment/depletion, and found that the collection of single amino acid substitutions selected in each screen are distinct (**Fig. 2D**). The first principal component of enrichment space explained 64% of the variation in the dataset, and largely separated the input library from the 48h screen, while the second principal component (17%) largely separated the 0h screen from the input. In the 0h screen, we characterized 96% (6,630) of single amino acid substitutions present in the SSV library in PAI-1 that met DESeq2-determined statistical thresholds as described in “*Experimental Procedures”* (**SI Fig. 1**). Of these, 3,376 single amino acid substitutions either maintained or improved PAI-1’s ability to inhibit uPA (log_2_-fold change > 0), while 3,254 variants resulted in reduced PAI-1 inhibitory activity toward uPA (log_2_-fold change) (**Fig. 2A**). For this screen we will henceforth refer to the log_2_-fold enrichment/depletion scores in our 0h screen as PAI-1’s *inhibition score*.

**Figure 2.**
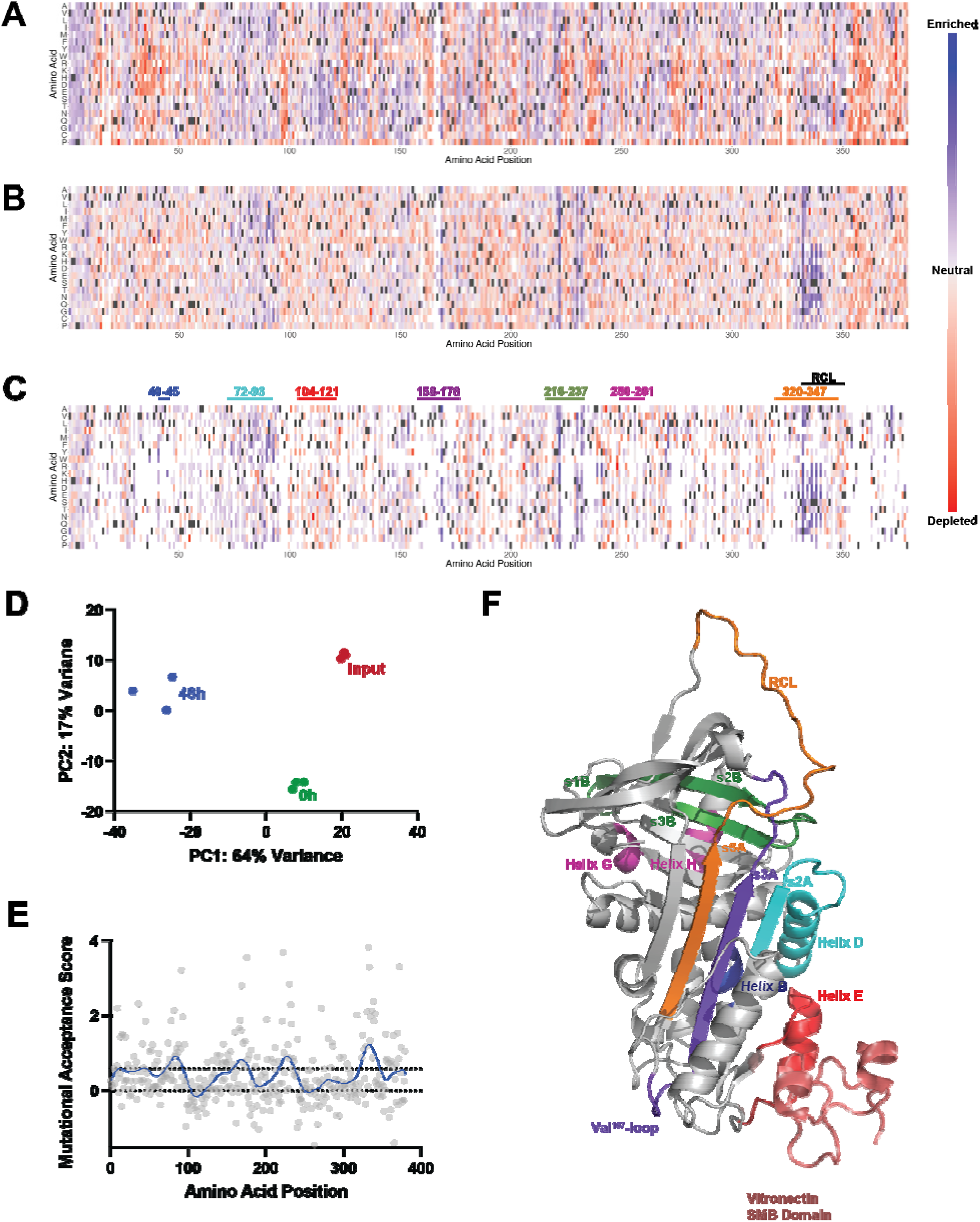
Sites of amino acid substitution that result in functional stabilization are clustered in flexible regions of PAI-1. (*A-C*) Heat maps of log_2_-fold change enrichment scores are shown for DMS screens following (*A*) 0h and (*B*) 48h incubations of the input variant library at 37 °C. Amino acid variants that were not present in the screened library are indicated in *white*. Panel (*C*) shows the results at 48h with the reduced function variants identified in the 0h screen (r*ed*) also indicated in *white*. Amino acid position is shown on the *x-axis,* while possible amino acid substitutions are shown on the *y-axis*. Canonical amino acids are shown in *grey* with enriched amino acid substitutions (log_2_-fold change > 0) in *blue* and depleted amino acid substitutions (log_2_-fold change < 0) in *red* relative to the input variant library. (*D*) PCA plot showing the first two principal components (PC1 and PC2) separating the input (*red*), 0h selected (*green*), and 48h selected (*blue*) libraries. (*E*) The log_2_-fold change results of the 48h screen corrected for non-function variants (*C*) were summed at each position (mutational acceptance score, *grey*), plotted as a function of amino acid position, and fit to a LOWESS regression (*blue*). Dotted lines indicate zero and the 75^th^ percentile of the LOWESS regression. (*F*) The AlphaFold2 (48) generated structure of PAI-1 (grey) is shown in the presence of the vitronectin SMB domain (PDB: 1OC0 (29), *red*). Regions in which amino acid substitutions functionally stabilize PAI-1 in its active conformation (as determined in *E* where the LOWESS regression was greater than the mean) are highlighted (as well as in *C*): residues 40-45 (helix B; *blue*), 72-93 (helix D and β-strand 2A; *cyan*), 158-178 (β-strand 3A and the Val^157^-loop, *purple*), 216-237 (β-strands 1B, 2B, and 3B; *green*), and 320-347 (β-strand 5A and the RCL; *orange*). Two regions in which amino acid substitutions tend to reduce the functional stability of PAI-1 (as determined in *E,* where the LOWESS regression was less than zero) are also highlighted, residues 104-121 (helix E which corresponds to the known binding site of vitronectin’s SMB domain; *red*) and 250-261 (helices G and H; *magenta*).

The library was next screened following a 48h incubation at 37°C to identify PAI-1 variants with either decreased or prolonged functional half-lives in PAI-1’s metastable active conformation. Following the 48h incubation of the 6,630 single amino acid substitutions characterized, 2,888 maintained their ability to inhibit uPA (log_2_-fold change > 0), while 3,742 displayed a reduced capacity to inhibit uPA (log_2_-fold change < 0, **Fig. 2B**). The later represents a combination of PAI-1 variants that had reduction inhibitory capacity toward uPA in the initial 0h screen as well as those with decreased functional stability; therefore, we next performed a post-hoc refinement of the results of our 48h selection to only include PAI-1 variants that at least maintain their uPA inhibiting capacity by removing those variants from further analysis where the inhibition score was less than zero in our initial 0h screen (**Fig. 2C**) (17). In this screen, we refer to the log_2_-fold enrichment/depletion score as PAI-1’s *functional stability score,* in which scores greater than zero represent PAI-1 variants with increased/maintained functional stability relative to wildtype (WT) PAI-1 and scores less than or equal to zero represent PAI-1 variants with decreased functional stability. This resulted in the identification of 3,376 single amino acid substitutions in PAI-1 that allow it to remain a uPA inhibitor but either prolong/maintain (functional stability score > 0, n = 2120) or reduce (functional stability score < 0, n = 1256) the rate of its latency transition (**Fig. 2C**). Of those that prolonged or maintained the rate of the latency transition, 331 single amino acid substitutions prolonged PAI-1’s functional stability beyond that of WT (variant functional stability score greater than that of the WT amino acid at a given position; **SI Data 2**).

Next, we confirmed if the PAI-1 regions in which single amino acid substitutions can prolong PAI-1’s functional half-life are the same as those identified previously (17). We averaged the functional stability scores at each amino acid position at PAI-1 to generate a *mutational acceptance score* with respect to prolonging PAI-1’s functional half-life and fit this score to a LOWESS regression (**Fig. 2E; SI Fig. 2**) (17). Regions in which the LOWESS regression was in its 75^th^ quartile were classified as functionally stabilizing, while those with a mutational acceptance score less than zero were classified as destabilizing (**Fig. 2C and 2F**). Using this approach, we identified five regions in PAI-1 where substitutions tend to increase PAI-1’s functional stability: residues 40-45 (helix A), 72-93 (helix D and β-sheet A strand 2), 158-178 (helix F, Val^157^-loop, and β-sheet A strand 3), 216-237 (β-sheet B strand 1-3), and 320-347 (β- sheet A strand 5 and the RCL (residues 331-350)). These regions grossly correspond to those identified previously in our lower-resolution DMS screen of PAI-1’s functional stability (**SI Fig. #3**) (17). The improved resolution of the screen presented herein, however, also allows us to identify the region between residues 105-121 (helix E) in which amino acid substitutions have the propensity to decrease PAI-1’s functional stability. This region encompasses part of the binding site for vitronectin’s somatomedin B (SMB) domain on PAI-1 (**Fig. 2F**) (29). As virtually all PAI-1 *in vivo* is found in complex with vitronectin, which modestly increases PAI-1’s functional half-life, this finding suggests that this region is not only critical for this macromolecular interaction but also for preserving PAI-1’s functional stability. This region is more than 40 Å from PAI-1’s RCL, suggesting individual amino acids in this region must function through long-range effects to stabilize or destabilize PAI-1 in its active conformation (30). The mechanism that mediates these long-range effects is unknown.

### Variant effect size prediction

To assess the correspondence between our DMS data and VEPs, we compared the results of our DMS screens for PAI-1 inhibitory function at 0h and 48h to three VEPs (EVE (23), CPT (22), and AlphaMissense (20) ; **Fig. 3**). All three VEPs differ in their methodologies and performance in benchmarking studies (21), with lower scores being indicative of a benign amino acid substitution and higher scores suggesting pathogenic amino acid substitutions. CPT was recently benchmarked (21) as the best predictor of PAI-1 inhibitory function among tested VEPs. However, given that all three VEPs performed similarly in these benchmarking studies, we anticipated that these predictors would all reflect the inhibition scores obtained in our present PAI-1 uPA inhibition screen.

**Figure 3.**
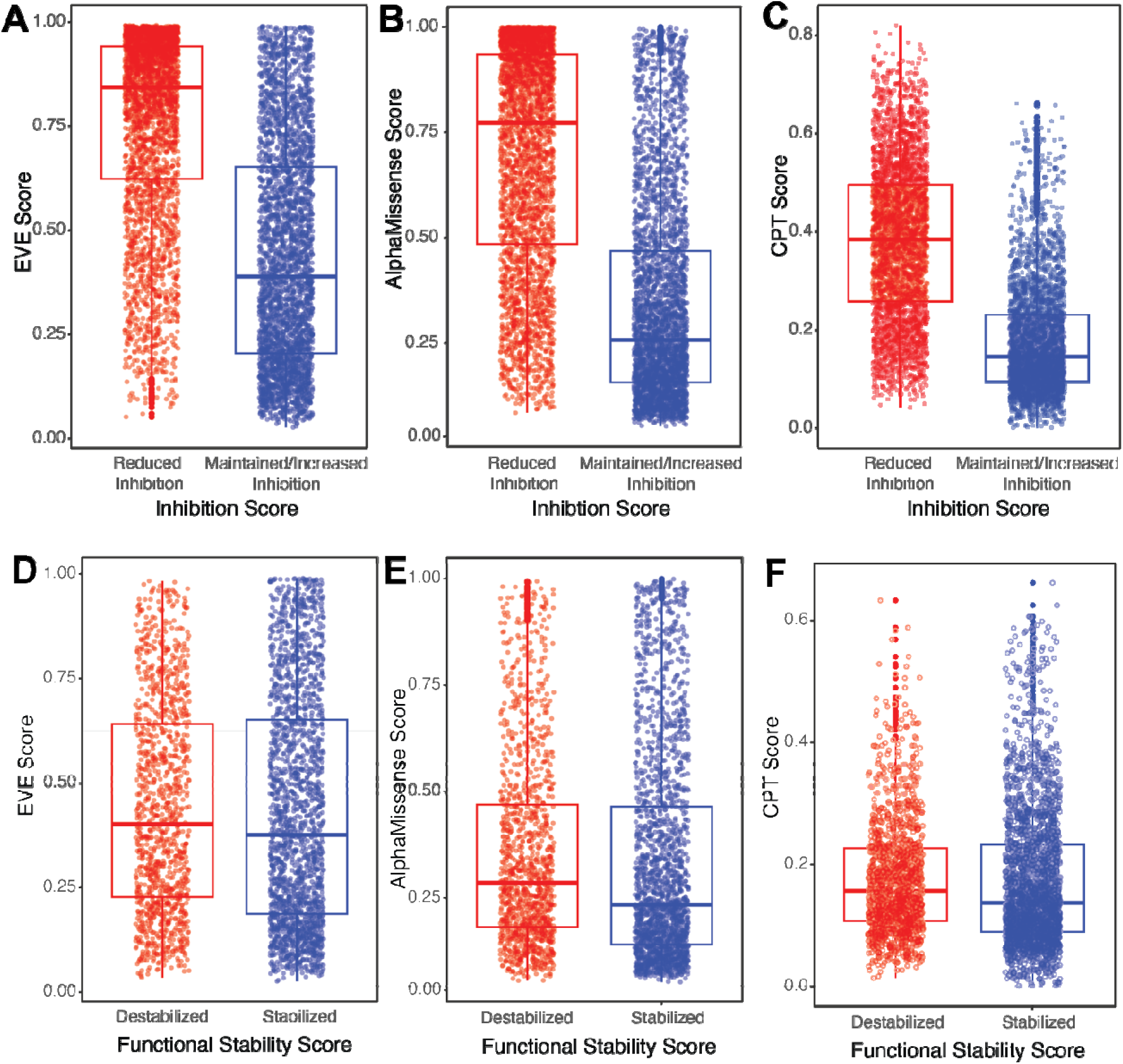
VEPs reflect the effect of amino acid substitutions on PAI-1 inhibition of uPA but not its latency transition. The VEPs (*A*) EVE, (*B*) AlphaMissense, and (*C*) CPT were compared to the results of the DMS screen for PAI-1 inhibition in which the inhibition score was stratified as either reduced inhibiton (inhibition score < 0, *red*) or maintained/increased inhibiotn (inhibition score > 0, *blue*). (*D*) EVE, (*E*) AlphaMissense, and (*F*) CPT were also compared to the results of the PAI-1 latency DMS screen in which the functional stability score was stratified as either destabilizing (functional stability score < 0, *red*) or stabilizing (functional stability score > 0, *blue*). The mean and 25^th^ and 75^th^ percentiles are indicated on each boxplot. The mean values and confidence intervals (CI), as well as the results of a linear mixed model analysis correcting for random positional effects are shown in **Table 1**.

**Table 1.**
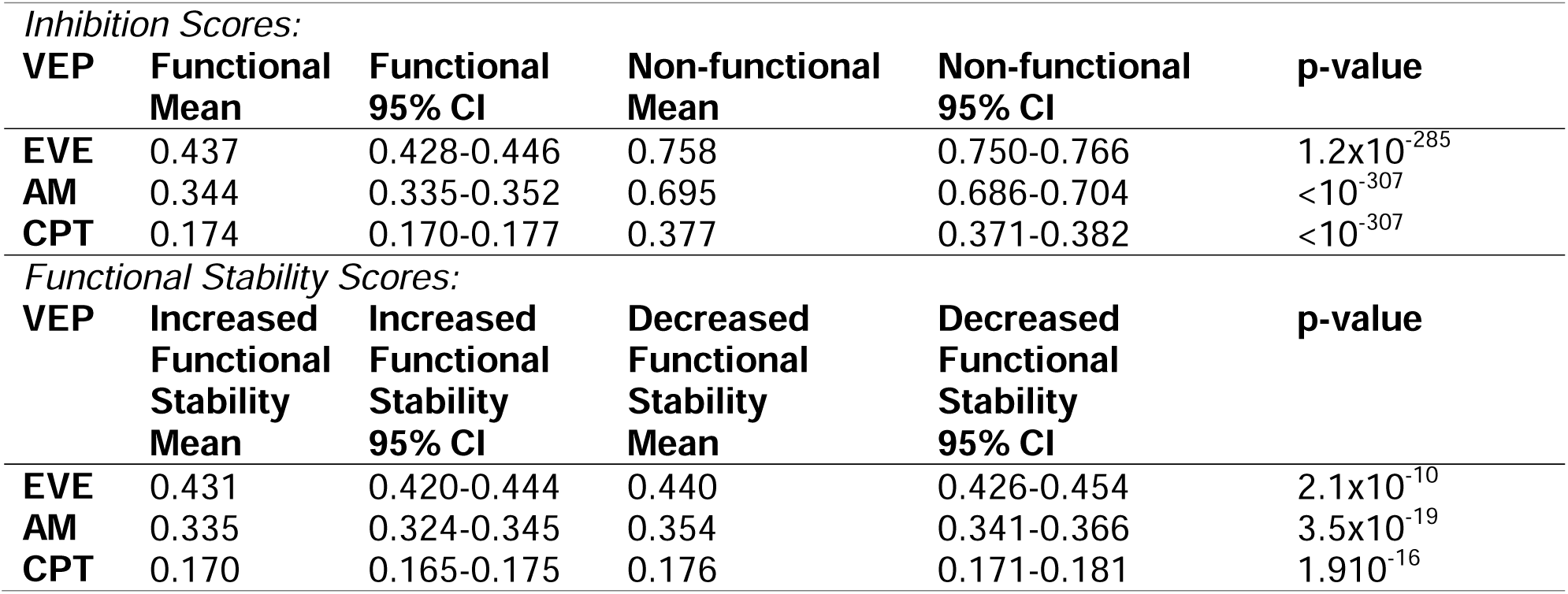
Summary of VEPs with respect to PAI-1 inhibition of uPA and latency transition.

| *Inhibition Scores:* |  |  |  |  |  |
| --- | --- | --- | --- | --- | --- |
| **VEP** | **Functional Mean** | **Functional 95% CI** | **Non-functional Mean** | **Non-functional 95% CI** | **p-value** |
| **EVE** | 0.437 | 0.428-0.446 | 0.758 | 0.750-0.766 | $1.2 \times 10^{-285}$ |
| **AM** | 0.344 | 0.335-0.352 | 0.695 | 0.686-0.704 | $<10^{-307}$ |
| **CPT** | 0.174 | 0.170-0.177 | 0.377 | 0.371-0.382 | $<10^{-307}$ |
| *Functional Stability Scores:* |  |  |  |  |  |
| **VEP** | **Increased Functional Stability Mean** | **Increased Functional Stability 95% CI** | **Decreased Functional Stability Mean** | **Decreased Functional Stability 95% CI** | **p-value** |
| **EVE** | 0.431 | 0.420-0.444 | 0.440 | 0.426-0.454 | $2.1 \times 10^{-10}$ |
| **AM** | 0.335 | 0.324-0.345 | 0.354 | 0.341-0.366 | $3.5 \times 10^{-19}$ |
| **CPT** | 0.170 | 0.165-0.175 | 0.176 | 0.171-0.181 | $1.910^{-16}$ |

To determine if the VEPs correlate with our empirical inhibition scores (from **Fig. 2A**), we stratified amino acid substitutions as either maintaining/improving (inhibition score > 0) or reducing (inhibition score < 0) PAI-1’s ability to inhibit uPA and compared the VEP score using a linear mixed model analysis correcting for random positional effects. As VEPs fill a need to identify disease causing or pathogentic mutations within the human genome, all three VEPs predicted that PAI-1 variants with decreased uPA inhibitor capacity were more likely to be pathogenic than those that were able to inhibit uPA with high statistical significance as anticipated (**Table 1**) (**Fig. 3A-C**). We next compared the results of our functional stability DMS screen (**Fig. 2C**) with the VEPs’ output on the basis of whether a substitution either *destabilizes* (functional stability score < 0) or *stabilizes* (functional stability score > 0) PAI-1’s in its metastable active conformation (**Fig. 3D-F**). All three VEPs predicted a slight, yet significant, increase in pathogenicity for PAI-1 variants with decreased functional stability (**Table 1**). Given that PAI-1’s latency transition is a conserved between fish and humans, we anticipated that amino acid substitutions that prolonged its functional half-life would be potentially pathogenic (15). However, these results suggest that these VEPs most readily predict the dominant function of PAI-1 as a uPA inhibitor but do not predict other collateral fitness effects, including PAI-1’s latency transition.

### Thermodynamics of the PAI-1 latency transition

To investigate the thermodynamics of the PAI-1 latency transition, we calculated ΔΔG values for single amino acid substitutions in PAI-1 using two algorithms, EvoEF (31) and FoldX (32). The ΔΔG values determined using these two methods are highly correlated with PAI-1 in both its active (PDB: 3Q02, chain B (33)) and latent (PDB: 1DVN (34)) conformations (**SI Fig. 4**). As described above in our VEP analysis, we first stratified inhibition scores as maintaining/increasing (inhibition score > 0) or decreasing (inhibition score < 0) PAI-1’s ability to inhibit uPA and compared their ΔΔG values in both the active (**Figs. 4A** and **4C**) and latent conformations (**Figs. 4B** and **4D**). Both EvoEF (**Fig. 4A-B**) and FoldX (**Fig. 4C-D**) predict that single amino acid substitutions in PAI-1 that retain its uPA inhibitory function are significantly more stable (lower ΔΔG) than those that render PAI-1 no longer capable of inhibiting uPA in both the active and latent conformations, consistent with the results of the VEP analyses (**Table 2**).

**Figure 4.**
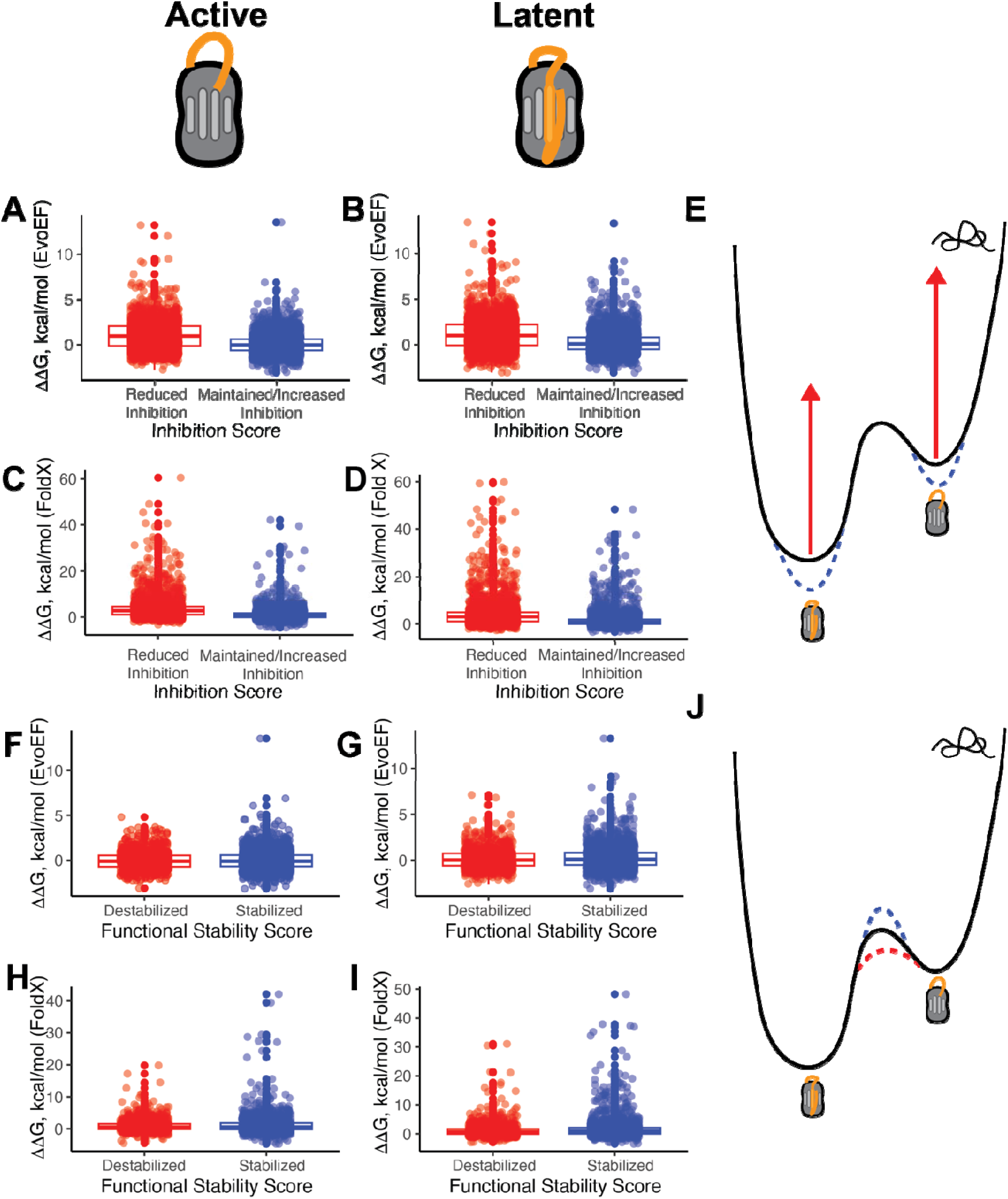
Thermodynamic stability predictions of PAI-1 variants in their active and latent conformations provide insight into the mechanism of PAI-1’s latency transition. (*A-D*) The results of the stability predictors (*A-B*) EvoEF and (*C-D*) FoldX were compared to the results of the DMS screen for PAI-1 inhibition in which the inhibition score was stratified as either reduced inhibition (inhibition score < 0, *red*, N = 3190 amino acid substitutions) or maintained/increased inhibition (inhibition score > 0, *blue*, N = 3253 amino acid substitutions) with PAI-1 in both its active (left column) and latent (right column) conformations. (*E*) A cartoon of the protein folding landscape highlighting PAI-1’s metastable active conformation and its lower-energy latent conformation illustrates that functional PAI-1 variants stabilize PAI-1 in both its active and latent conformations, while non-functional PAI-1 variants are most likely the result of protein misfolding. (*F-I*) The results of the stability predictors (*F-G*) EvoEF and (*H-I*) FoldX were compared to the results of the PAI-1 latency transition DMS screen in which the functional stability score was stratified as either destabilized (functional stability score ≤ 0, *red*, N = 1202 amino acid substitutions) or stabilized (functionality stability score > 0, *blue*, N = 1988 amino acid substitutions) with PAI-1 in both its active (left column) and latent (right column) conformations. (*J*) A cartoon of the protein folding landscape highlighting PAI-1’s metastable active conformation and its lower-energy latent formation illustrates that PAI-1 variants that tend to rapidly undergo a latency transition likely have a lower transition state energy for the latency transition (*red dashed line*), while PAI-1 variants that stabilize PAI-1in its active conformation are most likely the result of an increased energy barrier between the active and latent conformations (*blue dashed line*). The mean and 25^th^ and 75^th^ percentiles are indicated on each boxplot. The mean values and confidence intervals (CI), as well as the results of a linear mixed model analysis correcting for random positional effects are shown in **Table 2**.

**Table 2.** Summary of protein stability predictors (PSPs) with respect to PAI-1 inhibition of uPA and latency transition.

| *Inhibition Scores:* |  |  |  |  |  |
| --- | --- | --- | --- | --- | --- |
| **PSP** | **$\Delta\Delta G$**<br>**Functional**<br>**Mean**<br>**(kcal/mol)** | **$\Delta\Delta G$**<br>**Functional**<br>**95% CI**<br>**(kcal/mol)** | **$\Delta\Delta G$**<br>**Non-functional**<br>**Mean**<br>**(kcal/mol)** | **$\Delta\Delta G$**<br>**Non-functional**<br>**95% CI**<br>**(kcal/mol)** | **p-value** |
| **EvoEF**<br>**active** | 0.032 | -0.005-0.068 | 1.000 | 0.946-1.053 | $9.3 \times 10^{-59}$ |
| **FoldX**<br>**active** | 1.024 | 0.937-1.111 | 3.487 | 3.339-3.635 | $1.8 \times 10^{-59}$ |
| **EvoEF**<br>**latent** | 0.199 | 0.157-0.240 | 1.114 | 1.058-1.171 | $1.6 \times 10^{-49}$ |
| **FoldX**<br>**latent** | 1.420 | 1.309-1.531 | 3.978 | 3.797-4.160 | $1.0 \times 10^{-43}$ |
| *Functional Stability Scores:* |  |  |  |  |  |
| **PSP** | **$\Delta\Delta G$**<br>**Increased**<br>**Functional**<br>**Stability**<br>**Mean**<br>**(kcal/mol)** | **$\Delta\Delta G$**<br>**Increased**<br>**Functional**<br>**Stability**<br>**95% CI**<br>**(kcal/mol)** | **$\Delta\Delta G$**<br>**Decreased**<br>**Functional**<br>**Stability**<br>**Mean**<br>**(kcal/mol)** | **$\Delta\Delta G$**<br>**Decreased**<br>**Functional**<br>**Stability**<br>**95% CI**<br>**(kcal/mol)** | **p-value** |
| **EvoEF**<br>**active** | 0.024 | -0.024-0.072 | 0.011 | -0.046-0.067 | 0.61 |
| **FoldX**<br>**active** | 1.052 | 0.933-1.171 | 0.917 | 0.806-1.038 | 0.34 |
| **EvoEF**<br>**latent** | 0.228 | 0.173-0.283 | 0.138 | 0.074-0.201 | 0.31 |
| **FoldX**<br>**latent** | 1.581 | 1.426-1.738 | 1.112 | 0.970-1.254 | 0.83 |

This finding suggests PAI-1 variants that are incapable of inhibiting uPA trend toward being misfolded and, therefore, non-functional (**Fig. 4E**).

Next, we compared the results of our 48h functional stability DMS screen to ΔΔG calculations of PAI-1 in its active and latent conformations. Again, as described above for our VEP analysis, we stratified functional stability scores as either decreasing (*destabilized*) or maintaining/increasing functional stability (*stabilized*) (**Fig. 4F-I**). In both PAI-1’s active and latent conformation, there was no statistically significant difference in the ΔΔG values of PAI-1 variants with respect to their functional stability (**Figs. 4F-4I** and **Table 2**). This finding suggests that stabilization of PAI-1’s latent conformation does not significantly contribute to PAI-1’s latency transition. Rather, we speculate that in PAI-1 variants with increased functional stability, the activation energy to transition to the latent conformation increases, resulting in a kinetically slower process (**Fig. 4J**).

### Purifying selection of PAI-1’s latency transition and inhibitory function

In previous studies, we have shown that DMS with respect to uPA inhibition can be a powerful predictor of sites in PAI-1 that are evolutionarily conserved (27, 28). As the latency transition of PAI-1 is also reported to be an evolutionarily conserved trait of PAI-1 (15), we now expand our approach to this protein feature to understand the joint roles of latency and uPA inhibitory function in patterns of PAI-1 sequence evolutionary conservation. We hypothesize that the selection pressures on PAI-1 maintain both its latency transition and its primary function as a plasminogen activator inhibitor, and expect residues involved in both functions to be the most conserved.

For each position in PAI-1, we determined the *normalized functional mutation count* (the number of functional amino acid substitutions at each position (log_2_-fold change > 0 divided by the total number of scored mutations) with respect to both PAI-1 function as a uPA inhibitor, as well as its functional stability. If a trait is under purifying selection to conserve a function, then there should be a positive correlation between the normalized mutation count with respect to that trait and a conservation score determined by the degree of variation at the position across extant species’ PAI-1 orthologous sequences. In other words, if a trait is under purifying selection, the amino acid residues that are constrained or labile with respect to a given function will be so both across species and in our DMS assays.

We used ConSurf (35) scores based on PAI-1 sequences from 94 extant vertebrate species, where higher scores indicate more evolutionary lability (less residue conservation). We observe a significant positive correlation between the normalized functional mutation count (**Fig. 5A**; slope = 1.27, R^2^ = 0.155, p = 4.4x10^-15^), consistent with our previous data (27), suggesting that purifying selection maintains uPA inhibitory capacity. When we extend this analysis to PAI-1’s functional stability, we observe a negative correlation (**Fig. 5B**) in which amino acid residues in our screen that are most accepting of amino acid substitutions while remaining functional at 48h are more likely to be conserved in extant vertebrate species (slope = 0.73, R^2^ = 0.029, p = 1.3 x 10^-3^). Furthermore, with respect to both PAI-1 inhibitory function and functional stability, amino acid residues more accepting of amino acid substitutions are distributed throughout PAI-1’s primary sequence (**Figs. 5A-B**). These findings are consistent with selection to maintain PAI-1’s latency transition, albeit much less strongly, than selection for its canonical function as a uPA inhibitor.

**Figure 5.**
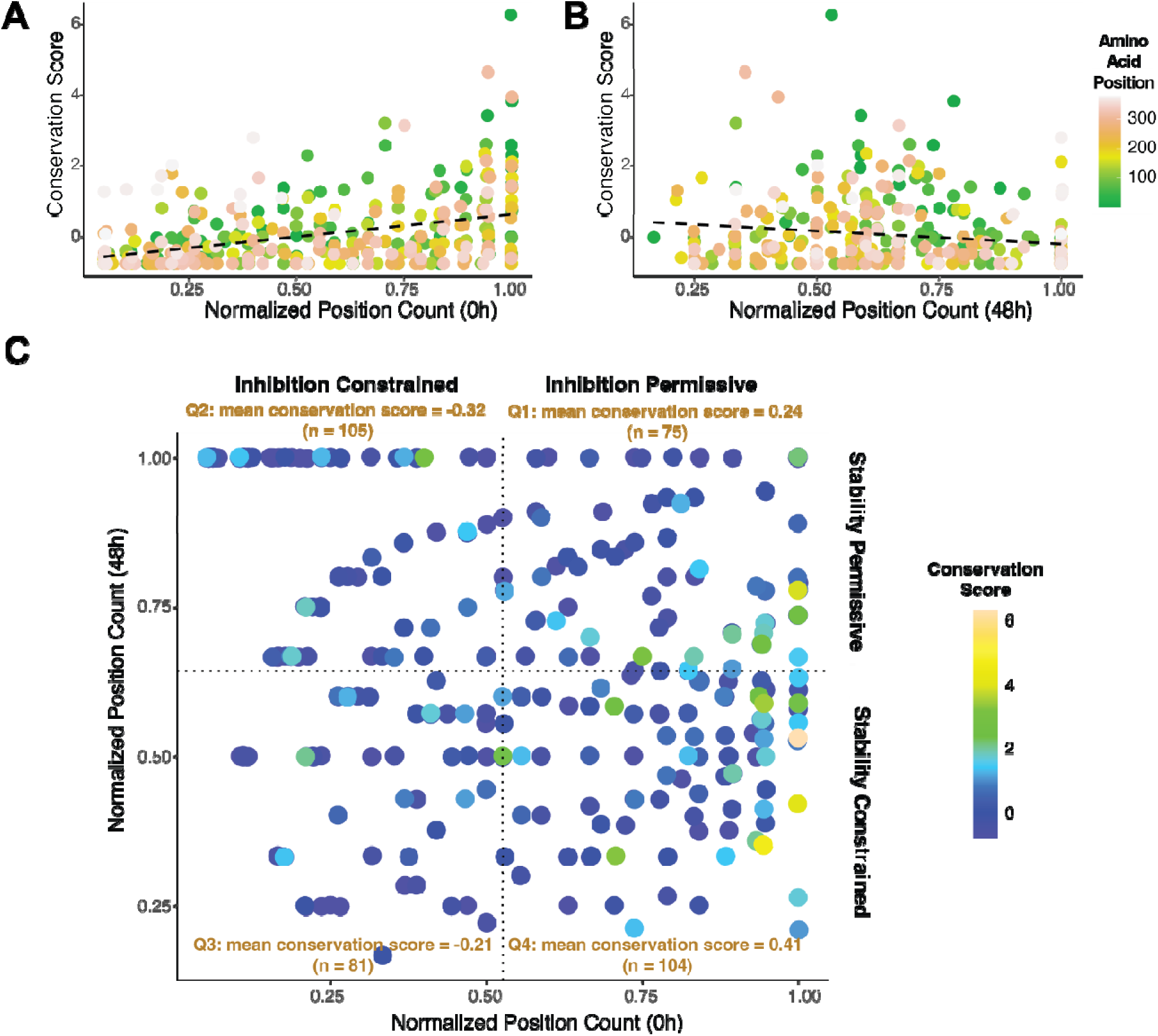
PAI-1’s inhibitory function and its latency transition are mutually evolving. (*A-B*) Conservation scores (35) for each amino acid position in PAI-1 are shown as a function of normalized position counts with respect to both (*A*) uPA inhibition (0h; slope = 1.3, R^2^ = 0.16, p = 4.9x10^-16^) and (*B*) functional stabilization in PAI-1’s active conformation (48h; slope = -0.73, R^2^ = 0.03, p = 0.0013). Amino acid position is colored on a gradient as indicated in the key. The best-fit linear regression is indicated by the *black* dashed line. (*C*) The normalized position count with respect to function (0h) is compared to the normalized position count with respect to functional stabilization (48h) for each variant. The conservation score is indicated by the color of the points, and the mean score is indicated in each quadrant. Tukey’s multiple comparisons test was used to determine statistically significant differences in the conservation score between quadrants (Q1 vs. Q2, p = 7x10^-4^; Q1 vs. Q3, p = 0.02; Q1 vs. Q4, p = 0.65; Q2 vs. Q3, p = 0.86; Q2 vs. Q4, p = 4x10^-7^; Q3 vs. Q4, p = 9.4x10^-5^).

We next wanted to determine if amino acid residues that are under joint constraint from both inhibitory and stability-associated functions show different degrees of evolutionary conservation relative to residues largely associated with a single function in our DMS screen. To this end, we analyzed the conservation scores at each amino acid position in PAI-1 as a joint function of the normalized position counts at both 0h and 48h (**Fig. 5C**). Segmenting this space into four quadrants allowed us to compare the interconnected evolutionary pressures on PAI-1’s function and latency transition (**Fig. 5C; SI Fig. 5**). Amino acid positions in quadrant (Q) 4, where amino acid substitutions are tolerated with respect to uPA inhibition (*inhibition permissive*) while being constrained due to impacts on the stability have the highest mean conservation score (0.41), consistent with a relaxation of selection to maintain both uPA inhibition and the latency transition at these sites. However, this level of conservation is not statistically different from that in Q1 (mean conservation score = 0.24; p = 0.65) where substitutions tended to increase stability.

However, the Q4 conservation scores are statistically increased compared to those in Q3 where substitutions tended to affect both uPA inhibition and the PAI-1 functional stability (mean conservation score = -0.32; p = 4x10^-7^). These results suggest that while both uPA inhibition and a relatively rapid latency transition are conserved features of PAI-1, uPA inhibition exerts stronger purifying selection than latency. The amino acid residues that affect PAI-1’s ability to inhibit uPA can also limit or promote its latency transition. We next performed a two-way ANOVA to test the interdependence of uPA inhibition and PAI-1’s latency transition on the evolutionary conservation score at each position in PAI-1. This analysis showed that while the ability to inhibit uPA has a highly significant impact on the evolutionary conservation score (F_1,361_ = 69.69, p = 1.5x10^-15^), the latency transition does not (F_1,361_ = 2.36, p = 0.13); however, the interaction between these two properties of PAI-1 is statistically significant (F_1,361_ = 7.67, p = 5.9x10^-3^), reinforcing the notion that the interaction between PAI-1’s inhibitor function and its latency transition are interdependent and both contribute to restricting PAI-1’s available amino acid sequence space.

## DISCUSSION

### PAI-1’s inhibitory function and latency transition are functionally linked

By comparing the results of our DMS screens for both PAI-1 inhibitory function against uPA and its latency transition to residue-level evolved variation among vertebrate species, we demonstrate that both features of PAI-1 are likely experiencing purifying selection (**Figs. 5A** and **5B**) and that the degree of conservation at a residue is a joint product of its roles in uPA inhibition and the latency transition (**Fig. 5C**). PAI-1’s function as an inhibitor of uPA and tPA is well-established (although only uPA inhibitory function was included in the present study as a marker of PAI-1 functionality), as both humans and mice deficient in PAI-1 present with increased bleeding and fibrinolytic activity (6, 7, 36–39). In contrast, the function of PAI-1’s latency transition remains unknown despite its presence across species as divergent as fish and humans (15). Latency, however, does not appear to be a requirement for SERPIN inhibition of tPA and uPA, as the most closely related SERPIN to PAI-1, protease nexin-1 (*SERPINE2*), does not undergo a latency transition yet still inhibits uPA and tPA (19, 40). Furthermore, the single amino acid substitution that most stabilizes PAI-1 in its active conformation is I91L (16, 17), which represents a reversion to the ancestral amino acid residue in eukaryotic SERPINs (41, 42). Together, these observations provide circumstantial evidence, which the present study confirms, that there is indeed selection to maintain PAI-1 variants that are both potent inhibitors of plasminogen activators and undergo a latency transition. Amino acid substitutions in PAI-1, both in across extant species and in our DMS screens, occur most frequently when it preserves both of these functions (**Fig. 5C**), thus minimizing collateral effects of a substitution that would result in a gain- or loss-of-function with respect to one and not the other (24).

### The vitronectin binding site is essential for PAI-1 stability even in the absence of vitronectin

Our data demonstrate that amino acid substitutions at the PAI-1:vitronectin interface can destabilize PAI-1 in the absence of vitronectin (Fig. **2C** and **2E-F**). Although the vitronectin binding site on PAI-1 is more than 40 Å from the RCL, vitronectin prolongs PAI-1’s half-life by 50 to 100% compared to the free SERPIN through an as-of-yet not fully understood allosteric mechanism (19, 29, 30). Vitronectin’s SMB domain binding region in PAI-1 appears to be in a structurally sensitive region across SERPINs for imbuing SERPINs with enhanced functional stability (2). For example, the SERPIN tengpin from thermophilic *Thermoanaerobacter tengcongensis*, which grows at an optimum temperature of 75 °C, is a SERPIN uniquely adapted to this extreme environment. Tengpin will undergo a latency transition, akin to that of PAI-1, in the absence of a 51 amino acid N-terminal segment that binds to the same region as vitronectin’s SMB domain. As neither shares structural or sequence similarity, this is suggestive of convergent evolution to address the biochemical problem of “stabilizing” a protein in a functional metastable conformation (2, 43).Therefore, given how critical this region is for modulating SERPIN functional stability, we postulate, as supported by our DMS data, that residues in this region may be adapted to provide modest functional stability to the latency- prone PAI-1 molecule through long-range interaction and that any perturbation here is likely to increase the rate of PAI-1’s latency transition.

### Thermodynamic and kinetic drivers of the PAI-1 latency transition

The results of our functional stability DMS screen (**Fig. 2C**) demonstrate that amino acid substitutions in some of the most conformationally flexible regions of PAI-1 stabilize its active conformation (**Fig. 2E-F**) (18, 44, 45), supporting the hypothesis that PAI-1’s instability in its metastable active conformation is driven predominantly by a reduced activation energy of the transition relative to other SERPINs. The rate-limiting step in PAI-1’s latency transition is the RCL moving over SERPIN’s “gate region” (**Fig. 2F**; β-sheet C strands 3 and 4) in order to insert into β-sheet A (45). Increasing the rigidity of PAI-1 will slow the rate of this transition and increase the number of PAI-1 molecules populating its metastable conformation, as partial unfolding of PAI-1 is required for it to undergo its latency transition (18, 45). Our data support a paradigm in which amino acid substitutions in PAI-1 affect its latency transition by modulating the activation energy required for the partial unfolding of PAI-1.

### Secondary functional properties may not be identified by VEPs

By their very nature, VEPs are trained to detect pathogenic or disease-causing missense mutations in proteins; however, this may not directly correlate with an understanding of protein function. Under physiologic conditions, compensatory mechanisms, including functional redundancy of proteins, may mitigate protein variant pathogenicity (46). Additionally, the data in the present study suggest another mechanism by which VEPs may not give an accurate picture of protein function. Both uPA inhibition and its latency transition are conserved functions of PAI-1 (15), and our study demonstrates that the amino acid residues that affect either function are linked with each other (**Fig. 5**). Our data clearly show that there are amino acid substitutions in PAI-1 that maintain PAI-1 inhibition of uPA while prolonging PAI-’s functional half-life that are likely selected against in nature (**Fig. 5B**). Such collateral fitness effects are a consequence of introducing amino acid substitutions into a protein that may result in a gain-of-function with respect to one protein property, while result in a loss-of-function with respect to another (24).

The VEPs used in this study (EVE, AlphaMissense, and CPT) were able to grossly capture PAI-1 function with respect to uPA inhibition (**Fig. 3A-C**). Consistent with our interpretation of the data, the majority of these latter single amino acid substitutions most likely result in misfolded proteins and a consequent loss of function.

Therefore, to better understand how single amino acid substitutions in PAI-1 affect both its inhibitory function and latency transitions, we employed two complementary approaches of conservation analysis (**Fig. 5**) and PSPs (**Fig. 4**). Using these approaches, we were able to determine that both of these two properties of PAI-1 are under purifying selection. When mutations at a given site were likely to affect both functions, conservation was highest, underscoring how multifarious selection pressures can align to result in the highest levels of protein sequence conservation. Furthermore, we confirm that predictions of protein stability (EvoEF and FoldX) grossly recapitulate the results of our study identifying PAI-1 variants with a reduction in inhibitory function as a result of protein misfolding (**Fig. 4A-D**). PSPs, however, were not able to predict changes PAI-1 functional stability when using a latent conformation of the protein, suggesting that the overall energy of the PAI-1 in its latent state has minimal effects on the tendency of a variant to undergo this transition. Rather, this supports a model in which the kinetics of escaping PAI-1’s active conformation local energy minimum is most likely the driving force behind the rate of the latency transition.

## CONCLUSIONS

Metastable proteins, including SERPINs, present a unique space for exploring the effects of amino acid substitutions on both stability and function. PAI-1 presents an ideal case for studying how individual amino acids affect these facets of protein behavior as it spontaneously undergoes a latency transition to its lowest energy conformation on experimental timescales.

This permitted us to explore how both PAI-1’s function and functional stability are linked, highlighting possible collateral fitness that should be considered when studying the effects of missense mutations on an isolated function of a protein. Therefore, this study supports a framework in which multiple facets of protein function are considered when predicting variant effects.

## EXPERIMENTAL PROCEDURES

### Deep mutational scanning

A SSV library with nearly equal representation of all possible single amino acid substitutions constructed on the WT PAI-1 backbone (**SI Fig. 1**) was purchased from Twist Bioscience (South San Francisco, CA) and cloned into a modified pAY-FE plasmid (Genebank #MW464120) as described previously (**SI Table 1**) (17). The cloned library was transformed into electrocompetent XL1 Blue MRF’ *E. coli* (Agilent Technologies). The PAI-1 SSV library was produced and uPA selection assays were performed as described previously (17, 27).

DNA coding for the PAI-1 displayed phage were sequenced over twelve 150 bp overlapping amplicons using the primers listed in **SI Table 1** (17, 27, 28). High-throughput sequencing data were analyzed using the DESeq2 software package (47). Significance thresholds for DESeq2 were set to a BaseMean score ≥ 50. MA plots showing the distribution of PAI-1 variants relative to these thresholds are shown in **SI Fig. 6**.

### Mutational Acceptance Score

Mutational acceptance scores were calculated for each position within PAI-1 with respect to functional variants in our 48h stability screen (**Fig. 2C**) by averaging the log_2_-fold functional stability scores at each position (17). The data were fit to a LOWESS regression with a 20-point smoothing window determined with the GraphPad Prism software package (v. 10.3.1). Regions of increased functional stability were identified where the LOWESS regression was greater than its mean value, while regions of decreased functional stability were identified as those where the LOWESS regression was less than zero.

### Protein stability prediction (**ΔΔ**G)

The effects of single amino acid substitutions were determined using both the EvoEF (31) and FoldX (32) methodologies.

### Variant effect size prediction

AlphaMissense (20) and CPT (22) PAI-1 variant datasets were obtained from public repositories. PAI-1 EVE scores were obtained from Debora Marks (Harvard University, Cambridge, MA) (23).

### Conservation scores

Evolutionary conservation scores were calculated at each position in PAI-1 using the ConSurf software package (https://consurf.tau.ac.il/) as previously described (27, 28, 35). The *normalized position count* at each position in PAI-1 was determined as the number of enriched amino acid substitutions (log2-fold change > 0) divided by the number of amino acid substitutions analyzed at each position (27). Conservation scores were analyzed with respect to the normalized position count using the R software package (version 4.3.3).

### Statistical analyses

Statistical analyses were performed using the R software package. To determine quartiles of constrained/permissive inhibition with respect to inhibition and latency were determined by dividing the data at the median values with respect to normalized position count at 0h and 48h.

## SUPPORTING INFORMATION

This article contains supporting information consisting of five supplemental figures and one supplemental table (Supp_Thermodynamics_and_Evolution.docx), as well as two datasets resulting from the DESeq2 analysis (*SI_Data_1.xlxs and SI_Data_2.xlxs*).

## DATA AVAILABILITY

Bioinformatics code and the associated data files required to execute it are available at https://github.com/hayneslm/PAI1_latency_evolultion_and_thermodynamics. Raw sequencing files and associated code may be accessed at https://doi.org/10.7302/bggn-2y63.

## FUNDING AND ADDITIONAL INFORMATION

This work was supported by the National Institutes of Health grant (R35-HL171421; D.G.). The content is solely the responsibility of the authors and does not necessarily represent the official views of the National Institutes of Health.

