## Supplemental Table and Figures for "Thermodynamics and selection of the plasminogen activator inhibitor-1 latency transition"

SI Table 1. DNA Primers

| Primer | 5’ → 3’ Sequence |
| --- | --- |
| Variant Library Cloning Primers | |
| TwistLib_S1 | AAAAAAAGGCGCGCCAGACACAAAACTCATCTCAGAAGAGGATCTGGGTG |
| TwistLib_AS1 | TAGTCACCACCACCTGCG |
| HTS Sequencing Primers | |
| PAI-1 Amplicon 1 For | NNNNNNGAACAAAAACTCATCTCAGAAGAGGATCTG |
| PAI-1 Amplicon 1 Rev | NNNNNNGGGTGAGAAAACCACGTTGCG |
| PAI-1 Amplicon 2 For | NNNNNNTTTCAGCAGGTGGCGCAG |
| PAI-1 Amplicon 2 Rev | NNNNNNCTTGTCATCAATCTTGAATCCCATAGCTGC |
| PAI-1 Amplicon 3 For | NNNNNNATGCTCCAGCTGACAACAGGA |
| PAI-1 Amplicon 3 Rev | NNNNNNTGTGGTGCTGATCTCATCCTTGTT |
| PAI-1 Amplicon 4 For | NNNNNNGCCCTCCGGCATCTGTACAAG |
| PAI-1 Amplicon 4 Rev | NNNNNNTTGCTTGACCGTGCTCCGG |
| PAI-1 Amplicon 5 For | NNNNNNAAGCTGGTCCAGGGCTTCATG |
| PAI-1 Amplicon 5 Rev | NNNNNNCCCAAGCAAGTTGCTGATCATACC |
| PAI-1 Amplicon 6 For | NNNNNNATTCATCATCAATGACTGGGTGAAGAC |
| PAI-1 Amplicon 6 Rev | NNNNNNCTGGAGTCGGGGAAGGGAG |
| PAI-1 Amplicon 7 For | NNNNNNAATGCCCTCTACTTCAACGGC |
| PAI-1 Amplicon 7 Rev | NNNNNNGGGCGTGGTGAACTCAGTATAG |
| PAI-1 Amplicon 8 For | NNNNNNCCCATGATGGCTCAGACCAAC |
| PAI-1 Amplicon 8 Rev | NNNNNNGGCAGAGAGAGGCACCTCT |
| PAI-1 Amplicon 9 For | NNNNNNAGCATGTTCATTGCTGCCCC |
| PAI-1 Amplicon 9 Rev | NNNNNNTTCAGTCTCCAGGGAGAACTTGG |
| PAI-1 Amplicon 10 For | NNNNNNCAGGCTGCCCCGCCT |
| PAI-1 Amplicon 10 Rev | NNNNNNACGTGGAGAGGCTCTTGGTC |
| PAI-1 Amplicon 11 For | NNNNNNAGACAGTTTCAGGCTGACTTCAC |
| PAI-1 Amplicon 11 Rev | NNNNNNGGGGGCCATGCGGGC |
| PAI-1 Amplicon 12 For | NNNNNNTCATCCACAGCTGTCATA |
| PAI-1 Amplicon 12 Rev | NNNNNNTGCGGCCGCTGAACCACCTCC |

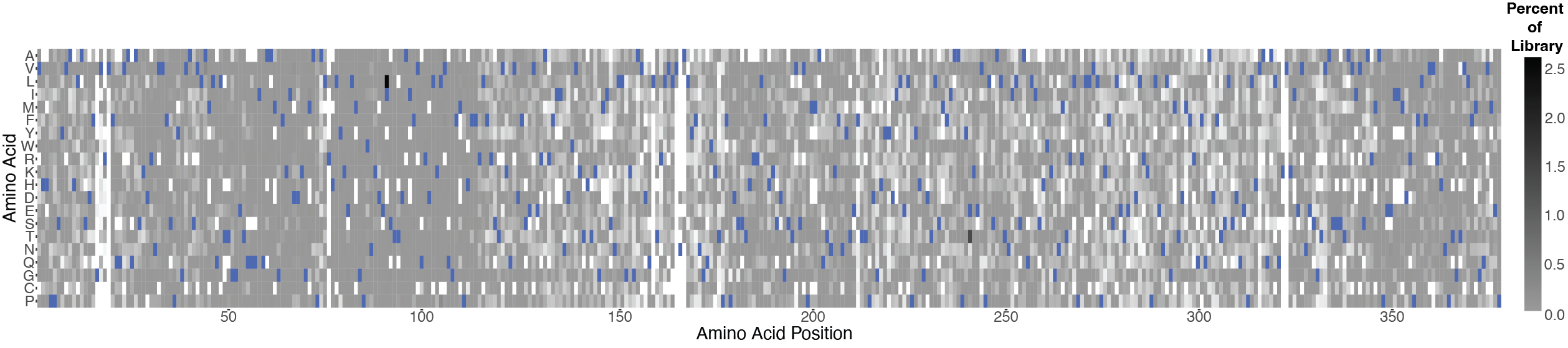

**SI Figure 1. Representation of single amino acid substitutions in the PAI-1 SSV library.** The PAI-1 SSV library used in this study contains 92% of all possible single amino acid substitutions. The amino acid position in PAI-1 is shown along the x-axis, with the possible amino acids along the y-axis. The fraction of the library represented by each variant in the input library as determined by Illumina sequencing is shown in greyscale, with the WT amino acids at each position shown in blue.

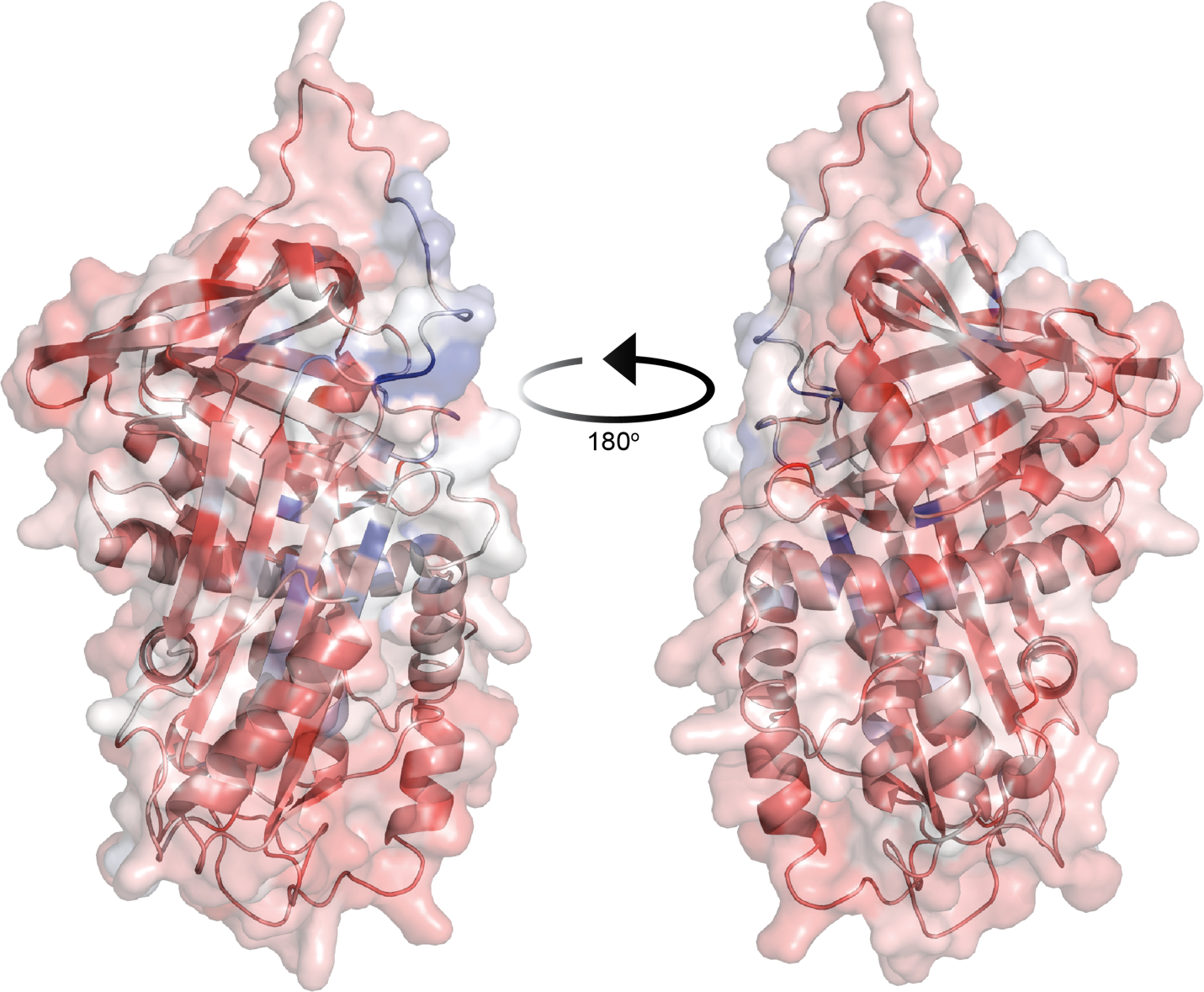

**SI Figure 2. Mutational acceptance scores mapped on the PAI-1 structure.** Mutational acceptance scores (Fig. 2E) were mapped onto the AlphaFold-derived structure of PAI-1 in its active conformation. Scores are mapped on a gradient of lowest (*blue*, most accepting of amino acid substitutions that stabilize PAI-1 in its active conformation) to highest (*red*, least accepting of amino acid substitutions that stabilize PAI-1 in its active conformation) with both frount (*left*) and rear (*right*) views shown.

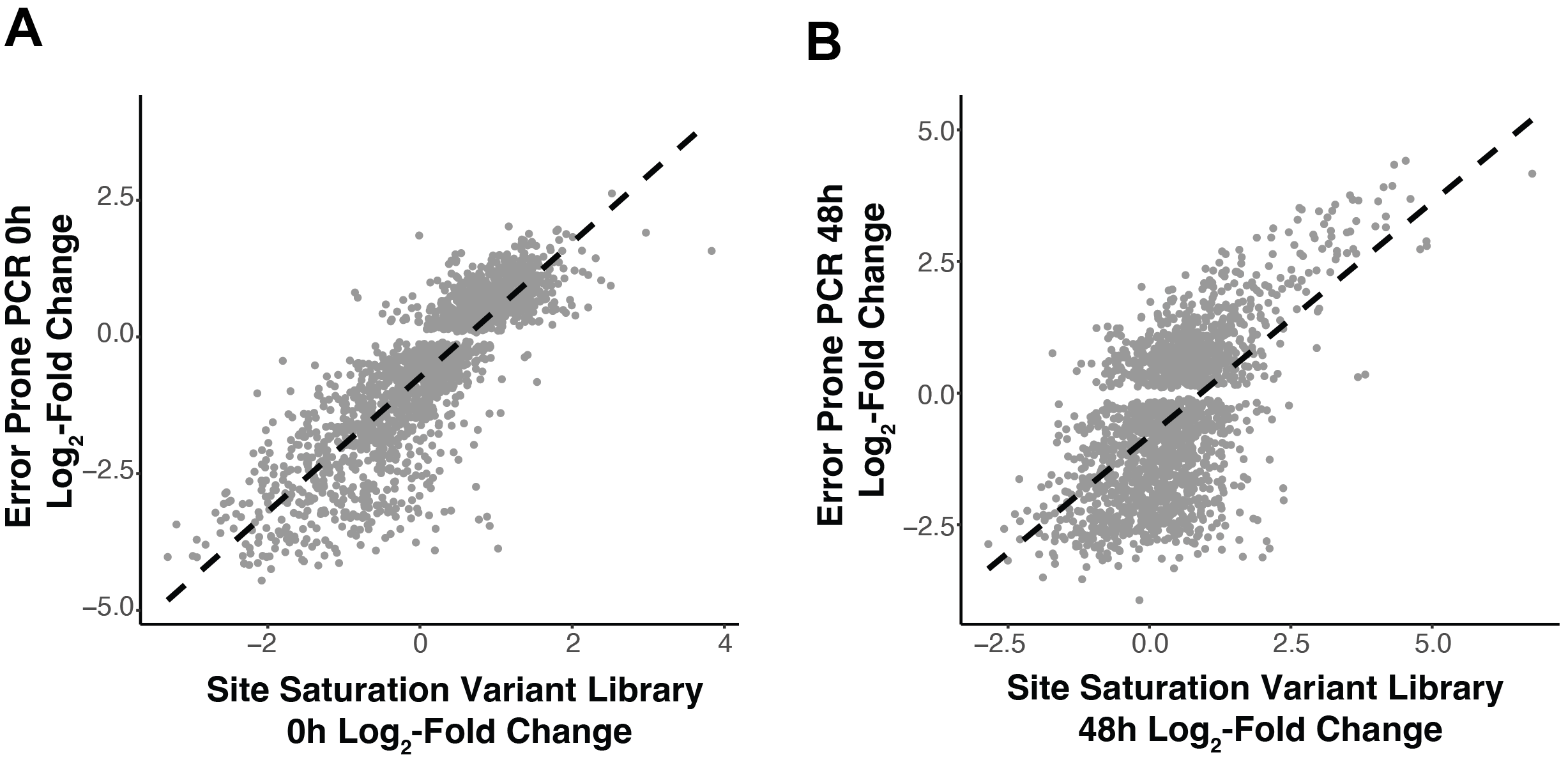

**SI Figure 3. Comparison of log_2_-fold changes scores in PAI-1 site saturation variant library and error prone PCR generated library.** (A) 0h screen comparing the PAI-1 site saturation variant library used in this study to those obtained using a variant library generated using error prone PCR (Huttinger, *et al*. *Sci. Rep.* 2021). A linear regression of the comparison is indicated by a dashed black line (slope = 1.23, R^2^ = 0.71, p < 2x10^-16^). (B) 48h screen comparing the PAI-1 site saturation variant library used in this study to those obtained using a variant library generated using error prone PCR (Haynes, *et al*. *JBC.* 2022). A linear regression of the comparison is indicated by a dashed black line (slope = 0.89, R^2^ = 0.36, p < 2x10^-16^).

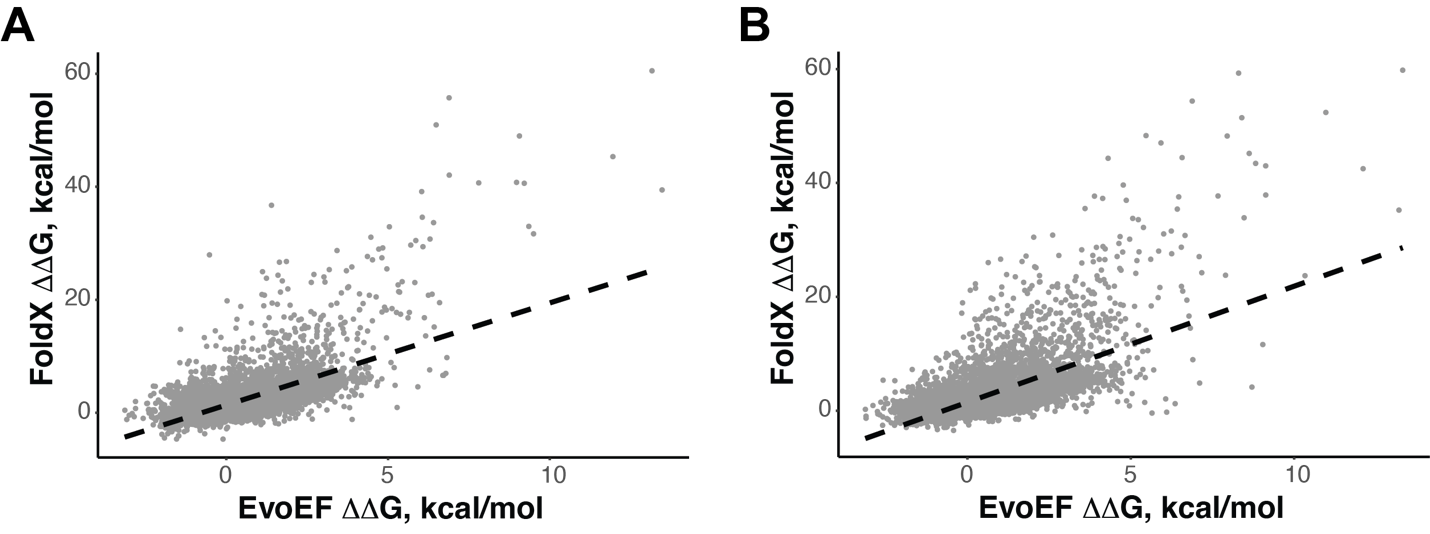

**SI Figure 4. Comparison of ΔΔG values determined using EvoEF and FoldX for PAI-1 in latent and active conformations.** ΔΔG values were determined for amino acid substitutions for PAI-1 in both its (A) latent (PDB: 1DVN) and (B) active (PDB: 3Q02) conformations. Individual amino acid substitutions are shown as grey points and the best-fit linear regression is shown as a dashed black line. The two models are moderately correlated in both the active (y-intercept = 1.5, slope = 2.0, R^2^ = 0.42, p < 2.2x10^-16^) and latent (y-intercept = 1.4, slope = 1.8, R^2^ = 0.42, p < 2.2x10^-16^) conformations.

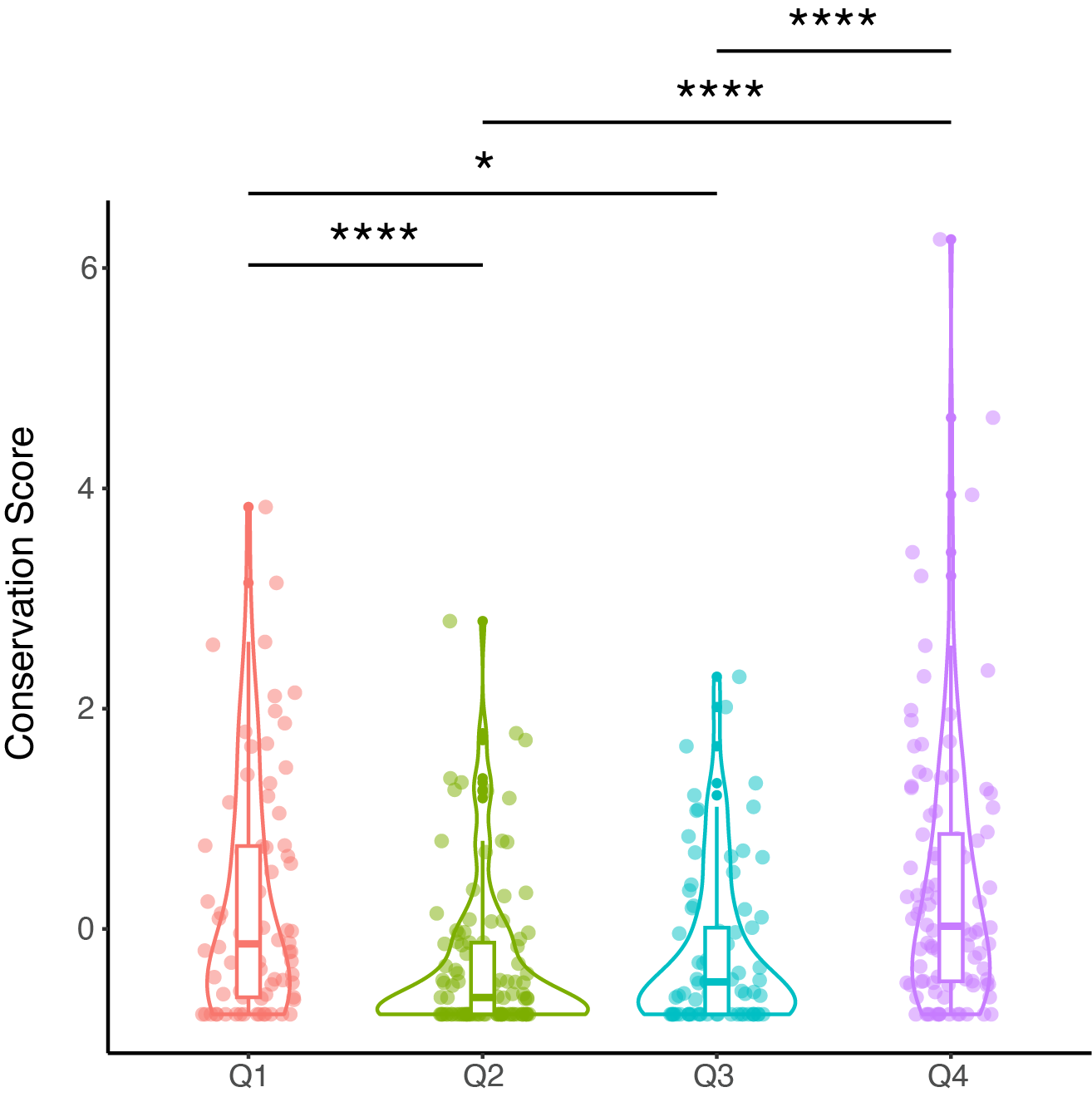

**SI Figure 5. Violin plot comparing quartiled conservation scores.** Conservation scores for each quartile (Q) defined in Fig. 5 are shown with each point representing a position in PAI-1. Statistical significance between the quartiles was determined using ANOVA analysis (*, p < 0.05; **, p < 0.01; ***, p <0.001; ****, p < 0.0001).

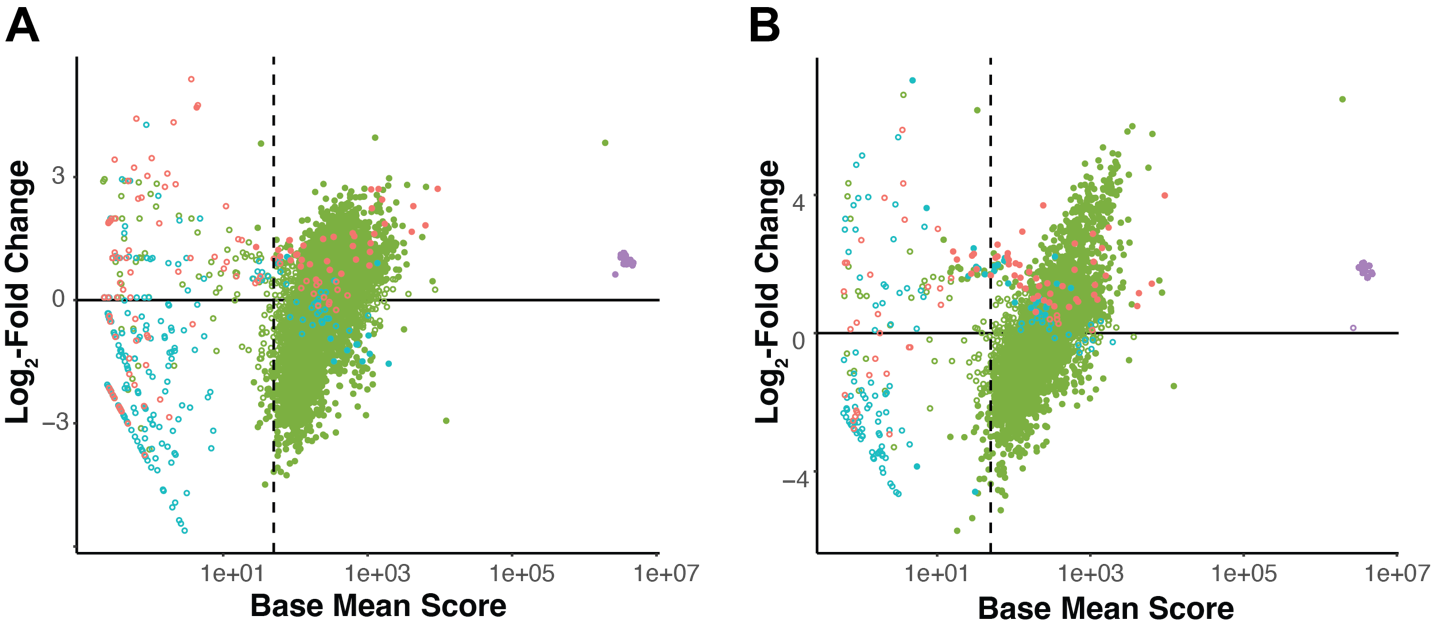

**SI Figure 6. MA plots for DMS screens of PAI-1 inhibition of uPA following 0h and 48h incubation at 37^o^C.** The log_2_-fold change in the enrichment/depletion scores for the SSV PAI-1 phage display library following (A) 0h and (B) 48h incubation at 37 ^o^C is shown as a function of the base mean score as determined by DESeq2 (Love, et al. 2014). Variants with a p_adj_ > 0.1 are shown as solid circles, and variants with p_adj_ ≤ 0.1 are shown as open circles. Colors indicate the type of mutation: WT amino acids (*purple*), missense mutations (*green*), amber stop codon (*salmon*), and all other nonsense mutations (*blue*).

**Description of Supplemental Data Files**

***SI_Data_1.xlxs*:** Results of DESeq2 analysis of the effects of single amino acid substitutions in PAI-1 on its ability to inhibit uPA following a 0h or 48h incubation of the phage displayed variant library at 37^o^C.

***SI_Data_2.xlxs*:** Results of DESeq2 analysis of the 48h dataset normalized for the log2-fold change enrichment score of the WT amino acid at each position (adj_score).
